# Disruption of Striatal Intrinsic Ignition in Huntington’s Disease: an In-Silico Perturbational Approach to Emulate Disease Progression and Recovery

**DOI:** 10.64898/2025.12.08.691368

**Authors:** Murat Demirtas, Dorian Pustina, Andrew Wood, Cristina Sampaio, Jakub Vohryzek, Gustavo Deco

## Abstract

Huntington’s disease (HD) is a neurodegenerative disorder caused by a single gene that severely affects motor, cognitive, and behavioral functions and is ultimately fatal. While disruptions in the structural integrity of the striatum are well documented, less is known about how the underlying functional impairments relate to the disease. We developed a large-scale modeling framework based on intrinsic ignition to uncover functional anomalies during HD progression. We first measured how much a spontaneous event in one brain region influences the rest of the brain, then systematically perturbed the activity of the striatum to predict the empirical changes in functional connectivity (FC) by re-optimizing the model parameters and conducting in-silico experiments. Results showed that disruption of the activity dynamics in the caudate, and to some extent the putamen, are the key factors that explain a substantial amount of the variance in FC alterations during HD progression. We then assessed the ability to recover whole-brain functional activity by driving the caudate and putamen activity towards a healthy-like regime and found that, by tuning the local bifurcation parameter of caudate and putamen, it is possible to recover whole-brain healthy functional activity, albeit moderately. Surprisingly, although the functional disruptions in the caudate better explained the alterations in FC, the recovery of the dynamics was more effective in the putamen. Overall, our results suggest that the disruptions of the dynamics in the striatum can explain the gross functional connectivity changes in HD. Our study demonstrates that computational modeling is an important tool to reveal the underlying patterns behind the observed data and provides an opportunity to test interventional mechanisms in silico.

## Introduction

Huntington’s disease (HD) is a debilitating neurodegenerative disorder characterized by motor, cognitive, and behavioral symptoms that typically appear between the ages of 40 and 50. Monogenic inheritance of HD is caused by an expanded CAG-repeat tract in exon 1 of the huntingtin (*HTT*) gene, which encodes polyglutamine. About 50 to 70% of the variation in age at symptom onset depends on the CAG-repeat length.

The neuropathological mechanisms underlying HD are not yet fully understood. However, substantial evidence indicates that the striatal structures—particularly the caudate and putamen—are severely affected, and medium spiny neurons in the striatum are lost at a higher rate than other neuron types; this degeneration is reflected in macroscopic volumetric atrophy measurable via magnetic resonance imaging (MRI) (Kinnunen et al., 2021). Atrophy in the caudate and putamen can be detected many years before clinical signs appear (Mena et al., 2023). This early detection has recently been leveraged as the best biological marker to detect the start of neurodegeneration well before the disease symptomatic stages develop, following the Huntington’s Disease Integrated Staging System (HD-ISS) (Tabrizi et al., 2022). The HD-ISS characterizes HD progression by distinct stages that span years before the emergence of clinical signs. Specifically, Stage 0 indicates individuals who carry the mutant *HTT* gene without any other signs, Stage 1 indicates biological degeneration reflected in the atrophy of caudate or putamen, Stage 2 indicates the emergence of the first clinical signs in motor or cognitive functions, and Stage 3 indicates loss of everyday functional abilities.

Characterizing early disease progression will be crucial to early therapeutic intervention. The HD-ISS categorization also raises questions about how the structural brain abnormalities seen in Stage 1 and Stage 2 are related to functional connectivity. Some studies suggest that there is no simple relationship between structural and functional impairments in HD (Minkova et al., 2016; Müller et al., 2016). Therefore, it is important to investigate the structure/function relationships during the progression of HD to elucidate disease mechanisms.

Several studies have identified anomalies in cortical and subcortical resting-state functional connectivity (rs-FC) in individuals with HD (for a review, see (Pini et al., 2020)). The impairments in rs-FC are most evident in the same subcortical areas that also show structural degeneration, specifically in the caudate and putamen (Paulsen et al., 2004). Disrupted connectivity in these regions correlates with CAG-repeat length and the severity of motor dysfunction, such as those measured by the Unified Huntington’s Disease Rating Scale (UHDRS), and can also predict the future onset of clinical signs (Unschuld et al., 2012; Wolf et al., 2008). Other research has revealed alterations not only in the striatum but across entire resting-state networks (RSNs) that involve multiple brain regions (Dumas et al., 2013; Werner et al., 2014). For example, early changes are observed in the sensory-motor network (SMN) before clinical motor deficits appear, with later alterations found in visual and attention networks (Pini et al., 2020). Prior studies also report significant topological changes in network organization (Gargouri et al., 2016; Harrington et al., 2015).

To date, only a few studies have focused on the dynamic aspects of FC (or time-varying FC). One study used clustering on dynamic FC matrices to construct meta-states that appear over time (Espinoza et al., 2019). By comparing the dwell times of these dynamic meta-states, they showed that HD patients spend more time in a specific state, thus exhibiting less dynamism than healthy controls. They also reported abnormal connectivity between the putamen and cortex. Another study investigated the dynamics of resting-state networks derived from independent component analysis (ICA) and found reduced FC in multiple resting-state networks in manifest HD compared to pre-HD participants (Aracil-Bolaños et al., 2022). Both studies showed reduced dynamic connectivity between the default mode network (DMN) and the salience network, as well as decreased connectivity between the DMN and the central executive network.

In 2017, Deco and KringelBach introduced the “intrinsic ignition framework” to assess how spontaneous local activation events influence overall brain activity (Deco and Kringelbach, 2017). The intrinsic ignition measure evaluates the relative significance of each brain region on the brain’s dynamic activity by examining how local events spread. This model involves reconstructing oscillatory signals in phase-space, enabling dynamic functional analysis. The framework has been effectively used to identify differences in activation patterns between various brain states, including wakefulness and sleep (Deco et al., 2019), meditation (Escrichs et al., 2019), and healthy aging (Escrichs et al., 2021).

Here, we aimed to use a large-scale computational model to study the effects of activity perturbations in brain areas. For this purpose, the most straightforward approach is to use Stuart-Landau equations (or simply the Hopf model), which provide a phenomenological description of activity in each brain region, where the system follows the dynamics generated by the normal form of a Hopf bifurcation (Deco et al., 2017a). Despite lacking biophysical details, the model elegantly displays one of the most important dynamics observed in neuronal activity: the interaction between noise-driven fluctuations and oscillatory activity (Figure 1A). Using this model, previous studies have shown that brain areas with high influence on the rest of the brain (i.e., hubs) exhibit higher values than those involved in local processing (Deco et al., 2017a). In the clinical context, (Demirtaş et al., 2017) demonstrated that alterations in FC during the progression of Alzheimer’s disease (AD) can be explained by a global decrease in the bifurcation parameter across all brain regions. Perturbation approaches have been previously studied theoretically and successfully used in many conditions (Deco et al., 2018; Fernandes et al., 2022; Jobst et al., 2021; Sanz Perl et al., 2021). In one study, external stimulation of nodes in the Hopf model was employed to reproduce whole-brain patterns during the switch between sleep and wakefulness (Deco et al., 2019). Similar methods have also been applied to model disruptions in whole-brain dynamics in Parkinson’s disease (Saenger et al., 2017) or stroke (Idesis et al., 2022). Therefore, this framework is an excellent candidate for studying the effects of disruptions in the striatum through in-silico manipulation of the system.

**Figure 1.**
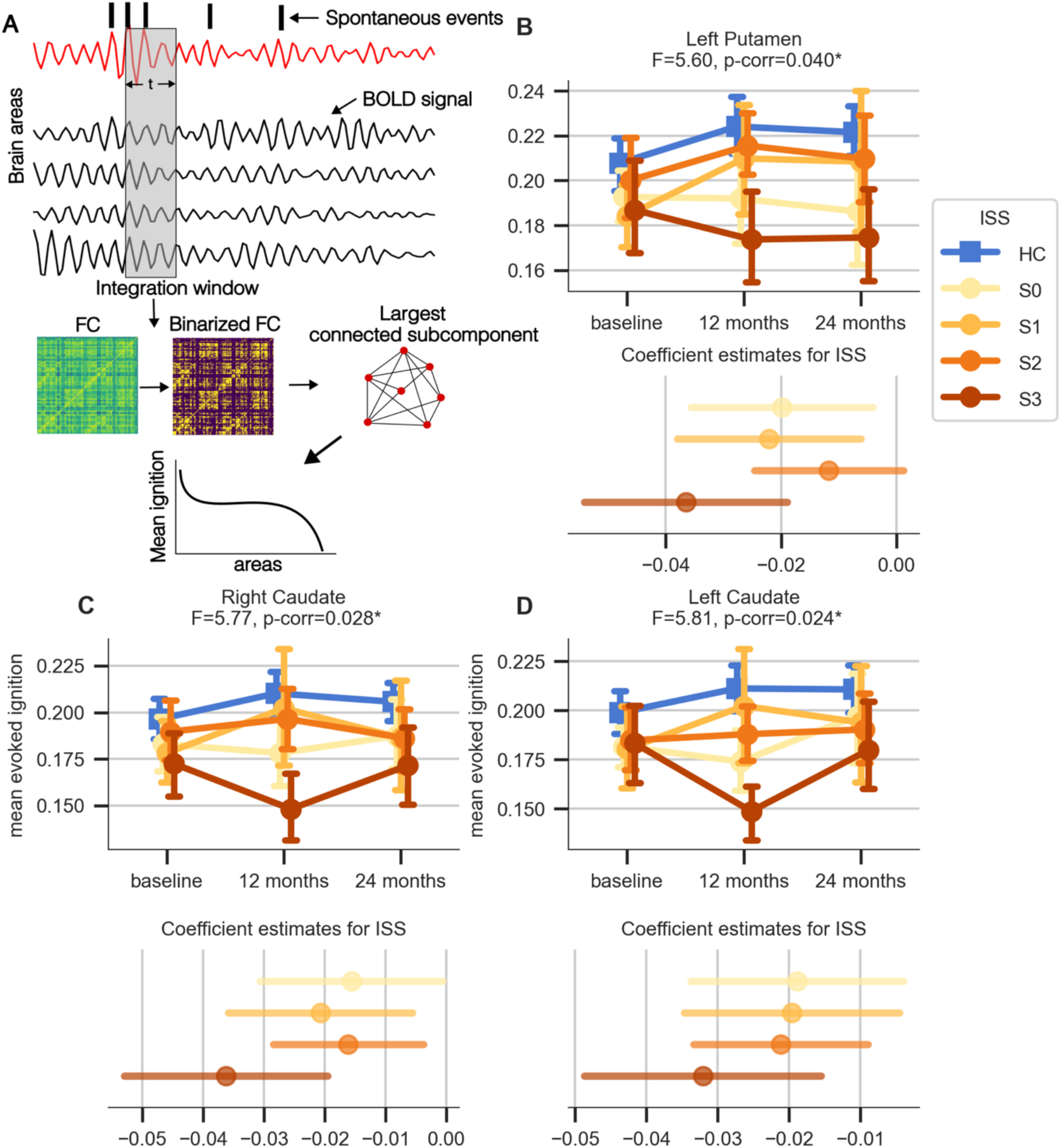
Linear mixed model results of average intrinsic ignition. **A** Schematic of intrinsic ignition framework. **B-D**. Results for left putamen **(B)**, right (**C**) and left (**D**) caudate. For each panel, upper part indicates average and standard error of each group/session, whereas the lower panel shows the coefficient estimates for each stage relative to HC modeled as a categorical variable. Only left putamen, left and right caudate are displayed because only those regions survived the max-F permutation correction, whereas right putamen still showed a statistical trend towards significance (Supplementary Materials).

In summary, we began this study by applying the intrinsic ignition framework to the HD dataset. We then inferred generative effective connectivity (GEC) matrices to establish the baseline in-silico dynamics. Next, we systematically perturbed the activity of the striatum to predict empirical changes in FC (Figure 1), which was done using three different approaches. First, we re-fitted the entire model by allowing the global bifurcation parameters and those of the caudate and putamen to vary freely. Second, we disrupted the dynamics of the caudate and putamen in models of healthy individuals to see if they replicate the FC changes observed across HD-ISS stages. Third, in people with HD (PwHD) models, we aimed to recover the dynamics of the caudate and putamen using another model that drives activity toward healthy-like dynamics.

## Material and Methods

### Study population

We used data from the TRACKOn-HD study (Tabrizi et al., 2011, 2009) harmonized in BIDS format and preprocessed with fMRIprep (Pustina et al., 2024). The data we used were from 240 participants with a maximum of three visits each. The gap between visits was 362,97 (STD: 67.13) days and 709.36 (STD: 58.77) days for session 2 and session 3, respectively. Of the 240 participants, 128 were HD gene carries and 112 of them were normative controls.

Following recommendations in the literature we removed scans with framewise displacement larger than 0.5 (Power et al., 2012). Upon inspection of the final timeseries we observed evident artifactual spikes in some subjects. To identify any artifactual spikes in the signal, we used absolute values of signal derivative. We checked the individuals in which the signal derivative above certain threshold and observed that for values above 6 standard deviations there are some shifts in the signal with unknown origin. If these deviations were observed during the first or the last 10 volumes, we excluded these volumes from the analyses and keep the session. However, if these deviations occurred in the middle of the recordings, it was not possible to remove these shifts due to short recording time of the data. Therefore, we excluded these individuals from the analyses. Of the remaining scans, 235 individuals had at least one rs-fMRI session.

No ISS classification could be made for 4 individuals; therefore, these subjects were excluded from the analyses. Among the remaining 231 individuals, there were 108 HCs, 21 STAGE 0, 13 STAGE 1, 60 STAGE 2 and 28 STAGE 3 individuals (Table 1).

### fMRI data acquisition and preprocessing

Data were collected at four imaging sites with similar acquisition parameters. The details of the data acquisition were provided in (Tabrizi et al., 2011, 2009). The data was acquired using a 3T Siemens Trio TIM, Philips Intera, and Philips Achieva MRI scanners depending on the site. Regardless of the scanner the resting state fMRI scans were collected using repetition time (TR) = 3000 ms, echo time (TE) = 30 ms, flip angle = 80, field of view = 64 x 60 x 48, voxel size = 3.131, total duration of 164 TRs.

fMRI data were preprocessed with the *fMRIprep* v20.2.7 pipeline (Esteban et al., 2019; Markiewicz et al., 2024); RRID:SCR_016216) during a data harmonization effort described in Pustina at al. (2024). We provide the full description of the processing steps in Supplementary Materials. In brief, the fMRI data were corrected for in-scanner motion, corrected for susceptibility-induced distortions using the acquired fieldmaps, band-pass filtered, co-registered to the T1-weighted structural scan, and resampled into several standard spaces including MNI152NLin2009cAsym, which was our space of reference for this study. Of note, we also applied the option for automatic removal of motion artifacts using independent component analysis (ICA-AROMA, Pruim et al. 2015) followed by spatial smoothing with an isotropic, Gaussian kernel of 6mm FWHM (full-width half-maximum). Corresponding “non-aggresively” denoised runs were produced after such smoothing. On the other hand, the “aggressive” noise-regressors were collected and placed in the corresponding confounds file. Several confounding time-series were calculated based on the *preprocessed BOLD*: framewise displacement (FD), DVARS, and three region-wise global signals. FD was computed following the Power et al. (2014) formulation of “absolute sum of relative motions”.

We used two parcellation templates to extract the fMRI signals from the timeseries: (1) the Schaefer-200 parcellation template for cortical regions (100 parcels per hemisphere) (Schaefer et al., 2018) and the Tian-16 template for subcortical regions (Tian et al., 2020). For convenience, we called this combined template as Schaefer-Tian parcellation in the rest of the manuscript.

### Structural connectivity

To estimate generative effective connectivity (GEC), we used a structural connectivity template derived from HCP dataset based on probabilistic tractography of diffusion weighted scans (DWI). The details of data acquisition are given in a previous study. (Cabral et al., 2022). In brief, the structural connectivity was derived from a probabilistic tractography-based normative connectome distributed with the leadDBS toolbox. This normative connectome was generated from diffusion-weighted and T2-weighted MRI data acquired in 32 healthy participants (mean age 31.5 ± 8.6 years, 14 females) from the Human Connectome Project. The diffusion-weighted MRI data were collected over 89 minutes on a specially designed MRI scanner with stronger gradients than those available on conventional systems. Diffusion data were processed using DSI Studio, applying a generalized q-sampling imaging reconstruction. A white-matter mask derived from the segmentation of the T2-weighted anatomical images was co-registered to the b0 volume of the diffusion data using SPM12. Within this white-matter mask, 200,000 most probable fibers were sampled for each participant. The resulting tractograms were then nonlinearly transformed to MNI standard space using a diffeomorphic registration algorithm based on deformation fields estimated from the T2-weighted images. Finally, the individual tractograms were aggregated in MNI space to generate a normative group tractogram representative of healthy young adults, which is provided within the leadDBS toolbox and from which the structural matrix was derived. We parcellated the resulting matrix using the Schaefer-Tian template.

### Intrinsic ignition framework

Intrinsic ignition characterizes the spatiotemporal propagation of information by measuring the degree of integration over time from spontaneously occurring events across the brain (Figure 2A). The framework is thoroughly described elsewhere in the literature (Deco and Kringelbach, 2017). In brief, we first obtained the ignition capacity of each brain region evoked by an event. The procedure of defining the events are adopted from Tagliazucchi et al. (2012). To define a binary event, we transformed timeseries into z-scores, *z_i_(t),* within a given time window (length of window = 3 TRs). Then we imposed a threshold, 8, such that the binary sequence of sequence σ(*t*) = 1 *if z_i_* (*t*) > θ and σ(*t*) = 0 otherwise. As we obtained the instantaneous phases of all brain regions, we calculated the phase lock matrix *P_jk_*(*t*), which characterizes the pair-wise phase synchronization between regions *j* and *k* at time *t* as follows:

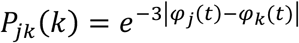

**Figure 2.**
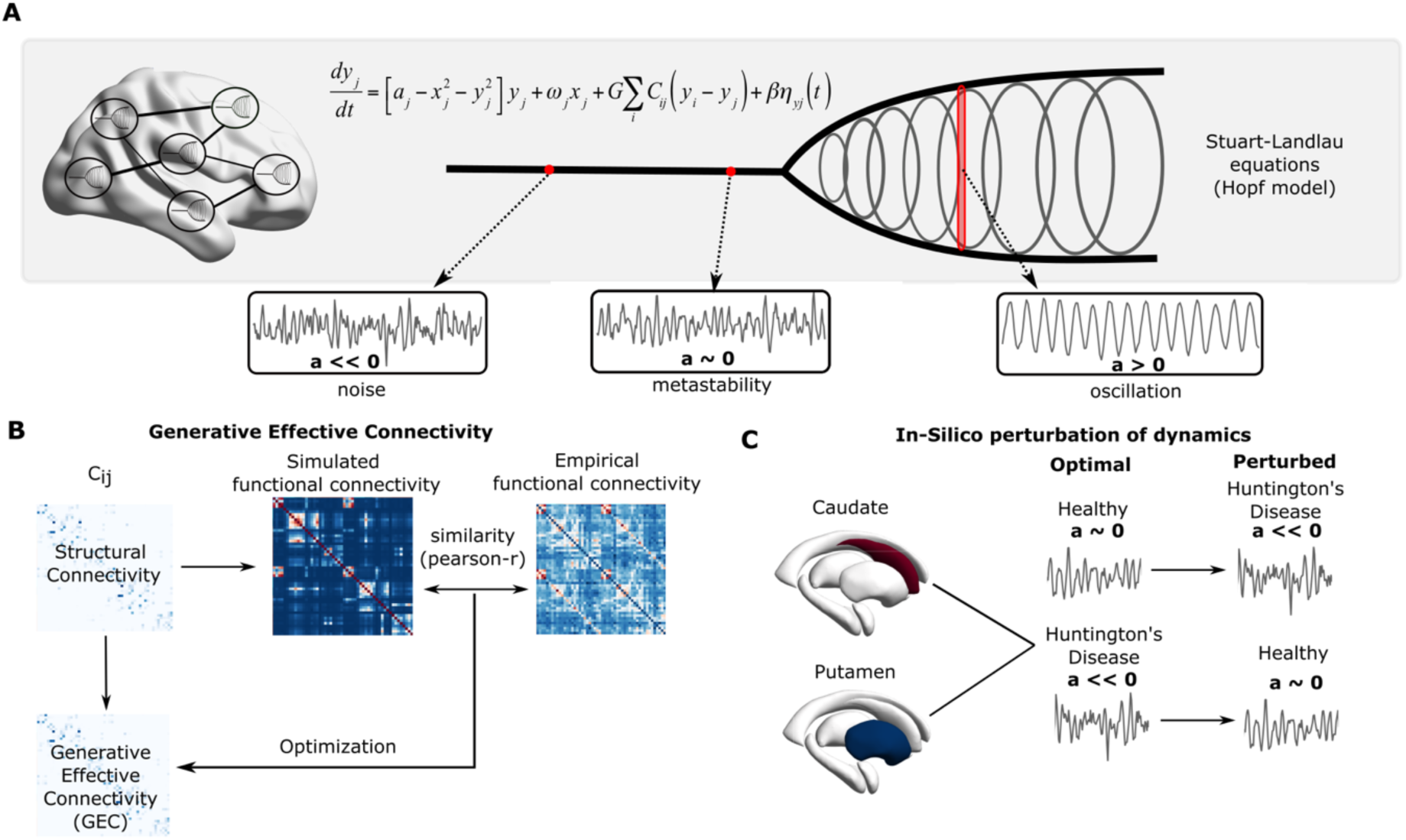
Schematic of the model framework. **A** Large-scale computational model. Each area is characterized by Stuart-Landlau equations, which is the normal form of a Hopf bifurcation. The local bifurcation parameter (a) exhibits Hopf bifurcation at value 0, that is the positive values produce oscillatory dynamics with the amplitude coupled to a, and negative values converges to a fixed point, hence producing noisy fluctuations. **B** Generative effective connectivity (GEC) estimation. Based on existing structural connections, GEC is estimated by minimizing the distance between empirical and simulated FCs. **C** In-silico computational experiments. Optimal bifurcation parameters of caudate and putamen is perturbed in a way that they are shifted towards noisy dynamics (i.e. a < optimal a) in healthy groups, and towards critical value (i.e. a ∼ 0) in HD groups.

Here, φ_j_ (t) and φ_k_ (t) indicate the phase of the brain regions *j* and *k* at time *t*. The integration is defined by the length of largest connected component in the binarized symmetric phase lock matrix *P_jk_*(*t*). We used a fixed threshold, θ, to binarize the phase-lock matrix such that (*C_jk_* = 0 if |*P_jk_*| < θ, 1 otherwise). The integration value is computed as the largest subcomponent, that is the length of the connected component in an adjacent graph, which measures the broadness of communication across the network for each driving event (Deco et al., 2015). Finally, we repeated this procedure for each event, each brain area, and computed the average integration for each brain region.

### Statistical analyses

We used linear mixed model (LMM) to make statistical inferences on mean integration acquired from intrinsic ignition framework. We used multiple models to identify the best fit. The first, most basic model was:

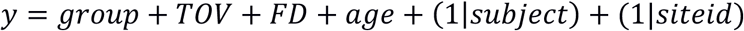

where y is mean intrinsic ignition, group is the assignment variable (group = {HC, STAGE 0, STAGE 1, STAGE 2, STAGE 3}, indicating healthy controls, STAGE 0, STAGE 1, STAGE 2 and STAGE 3 groups, respectively), TOV indicates the time of visit (baseline, 12-months and 24-months), while we controlled for age and framewise-displacement (FD). To model longitudinal data, we included subject intercept as a random effect in the model. Finally, we included site ID intercept as another random effect to take into account shared variance due to different recording sites.

The alternative model we considered also included the interaction between group and study TOV variables to test longitudinal effects:

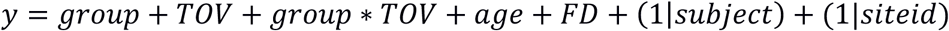

We compared the alternative model with each other using likelihood ratio test where applicable. Unless the alternative models explained the variance significantly better than the simpler one, we used the simplest model without interaction term and random slope. Finally, we summarized the statistics for the estimates of interest using F-statistic and its associated p-value. Where applicable, to illustrate the effect of progression across ISS stages, we also computed the relative coefficients (i.e. for each ISS stage relative to HC) considering groups as a categorical variable. To corrected for multiple comparisons across different areas, we fit the model after shuffling ISS and FD of the subjects and recorded the maximum F-statistic (ANOVA over the linear model) over 216 areas for 1000 permutations each (Winkler et al., 2016).

To compare the means of optimal model parameters across groups we used a non-parametric Kruskal-Wallis test, since the distributions did not appear normally distributed. Finally, to test whether the improvements in model fits were greater than 0, we use one-sided T-test.

### Generative effective connectivity model

The model contained 216 nodes, where each node was coupled with each other via DWI-derived structural connectivity (SC) matrix. The local dynamics of each individual node were represented by normal form of a supercritical Hopf bifurcation (Deco et al., 2017a, 2023; Kringelbach et al., 2023). The BOLD signals were described by Stuart-Landau oscillators:

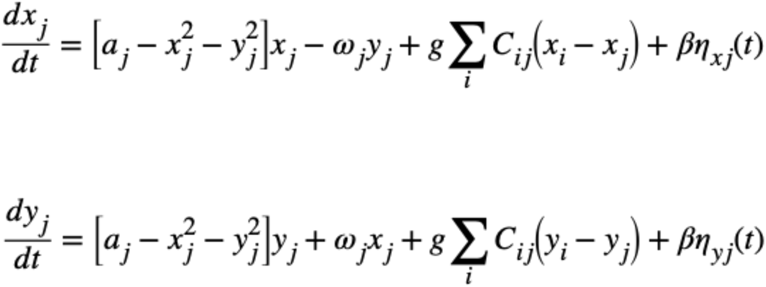

Where ω is the intrinsic frequency of each node, a is the local bifurcation parameter, η is additive Gaussian noise with standard deviation β, the temporal evolution of the activity, z, in node j is given in complex domain comprising x and y components, *C*_*ij*_is the Structural/Effective Connectivity between nodes i and j, g is the global coupling factor, and the standard deviation of Gaussian noise, β = 0.02. The natural frequency (f) of each region is taken as the peak frequency in the given narrowband of the corresponding region in the empirical time-series.

Following the approach previously employed (Deco et al., 2014), we analytically estimated the model FC using linearization of the system around a stable fix point. This approach offers a very fast way to calculate model FC, hence it is an efficient approach to estimate whole-brain generative effective connectivity (GEC). In brief, using the Taylor expansion of the system, the fluctuations around the fix point can be described as:

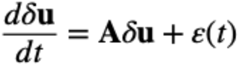

Where **A** is the Jacobian matrix, and e(t) is the noise term. The Jacobian matrix of the system evaluated at the fixed point can be constructed as:

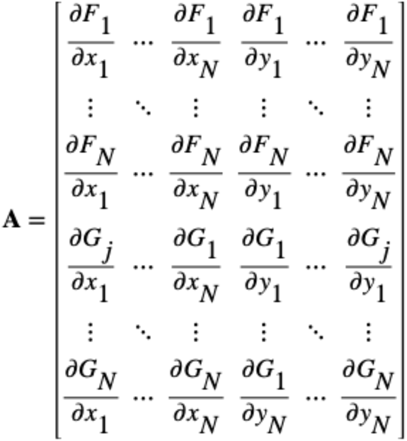

Then, each element in matrix A can be calculated as:

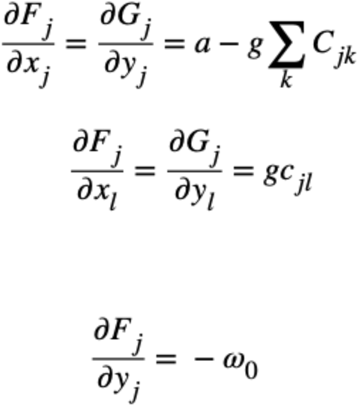

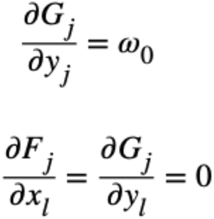

Where Q is the noise covariance matrix, the covariance matrix of the system P can be estimated by solving Lyapunov equation. Finally, the model correlation matrix (FC) can be extracted from the covariance matrix (P).

To find the optimal GEC, we start with a binarized version of the empirical structural connectivity matrix, which is used for identifying directly connected areas ignoring the actual anatomical connectivity strength. At each iteration, we assess the similarity between empirical and model FCs, while adjusting the underlying EC. Then we use a heuristic approach to find the optimal GEC matrix that maximizes the similarity between empirical and model FCs. Therefore, instead of using FC that reflects the involvement of many complex dynamic processes such as interactions across local oscillatory activity, noise, global signal and various artifacts, we search for a connectivity matrix (i.e. GEC) that can generate an approximation of FC with the constraints from SC matrix (i.e. no connections can be present between two nodes if there is no structural connectivity).

The model-generated functional connectivity (*FC^model^*) and time-shifted functional connectivity (*FC^tau,model^*) were used to infer the optimal generative effective connectivity (GEC) matrix that explains the observed functional connectivity (*FC^emprical^*) and time-shifted functional connectivity (*FC^tau,emprical^*). Tau for time-shifted FC is set to 1 TR (3 seconds). We estimated optimization procedure was performed as follows: First, we defined the error function as the sum of squared differences between model and observed FCs for both 0-time lag and time-shifted FC matrices:

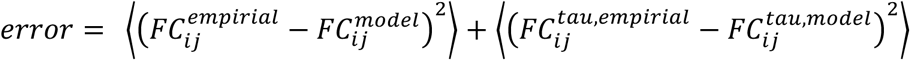

Where the initial guess for GEC (*C_ij_*) is the average SC matrix driven from Human Connectome Project (HCP). In GEC approach, SC only serves as a template, since the magnitudes of the connections are not relevant. Therefore, it is a common practice to use an average healthy template whenever the empirical DWI matrices are not available. The new guess for GEC is updated according to the equation:

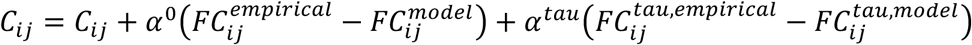

Here, ∝^0^ for FC is taken 0.0004 and ∝*^tau^* for FC-tau is taken 0.0001, t At each iteration, we set all the connections that are absent in SC to 0. This is aimed to avoid over-fitting by artificially assigning connectivity weights between brain areas that are not physically connected to each other (*C_ij_* = 0 *if SC_ij_* = 0). For convenience, to avoid overflow of GEC values, we also normalized the GEC between 0 and 0.2 (*C_ij_* = *C_max_C_ij_*⁄max (*C*)).

The iterations were repeated until the normalized error is lower than 0.001 or the error is no longer improving as defined by following equation:

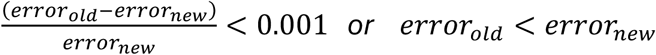

Or the maximum number of iterations (5000) is reached. During the optimization procedure none of the individuals reached the maximum number of iterations and all successfully converged to an optimal solution. This indicates that the optimization procedure was successful.

### *In-silico* Perturbation strategies

As described earlier, we fixed the bifurcation parameter a at −0.05 globally, while estimate GEC matrix of each individual. Based on these optimal solutions, first, we re-optimized the global, caudate and putamen bifurcation parameters. For this, we kept estimated GEC matrices and all other parameters the same. During the optimization, we defined the global bifurcation parameters (*a_global_*) brain regions other than caudate (*a_caudate_*) and putamen (*a_putamen_*) as homogeneous. Then, we optimized the similarity between model and empirical FCs, setting *a_global_* and *a_caudate_* or *a_putamen_* as free parameters. We used differential evolution (DE) to optimize the model (Storn and Price, 1997). Since DE algorithm is not guaranteed to find the global optimal solution, we run the algorithm 10 times for each individual and used the best fit among all optimizations.

### Experiment 1: Perturbing HC individuals towards HD state

As the empirical observation, we calculated the difference between average FCs of HC and those of each ISS stage:

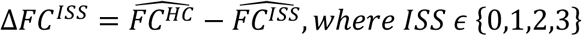

Then, using the average EC matrices of HCs, we calculated the baseline FC (unperturbed). Keeping all other parameters the same, we modified local bifurcation parameter of each brain area (*a_i_*) such that 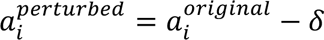. We fixed *δ* = 25 for convenience since the correlation values increased by the intensity of the perturbation but then stabilized, so we chose an extreme value to ensure stabilization. After calculating the FC with the perturbed values, we produced an in-silico counterpart for the FC difference:

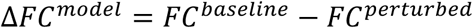

Finally, we calculated the correlation coefficient between Δ*FC^ISS^* and Δ*FC^model^* to quantify the similarity between empirically observed FC alterations and the predictions by the model.

After computing the values in average FC matrices, we repeated the procedure for each individual HC. For this, we simply estimated the same quantity 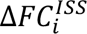 for each individual, *i*, and similarly calculated the simulated 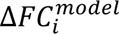 for each individual and then calculated the correlations between those values.

### Experiment 2: Recovering HD FCs in model

As the inverse of the same procedure provides very similar information, we proposed a slightly different experiment to recover HD FCs in the model. This time we used the native ECs and re-optimized the bifurcation parameter (for caudate and putamen) as the baseline model FC for each HD patient. Then we computed the correlation between the baseline model FC and average FC of the HCs (*r*^*baseline*^). We perturbed the optimal bifurcation parameter of each individual such that 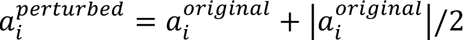. In other words, this perturbation pushed the bifurcation parameters of caudate and putamen halfway closer towards 0 (i.e. emulating the values closer to healthy individuals). Then, we computed the updated correlation between the perturbed model and the same average FC of the HCs (*r^perturbed^*). Finally, for each HD individual, we calculated percentage of the change in fit:

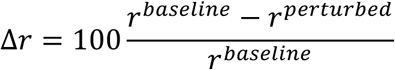

## Results

### Intrinsic ignition framework

First, we computed the average evoked ignition of each brain area for each individual. Then we constructed a linear mixed model to predict the observed values based on group category (HC, HD-ISS Stage 0, 1, 2, 3) and time of visit controlled for framewise displacement and age. After controlling for multiple comparisons using the permutation approach, we did not find any longitudinal effects. However, we found that intrinsic ignitions in bilateral caudate and left putamen were significantly reduced across the HD groups (Figure 1B-D). Overall, these results confirmed previous studies which found that the BOLD activity in striatum is disrupted in HD (Pini et al., 2020; Unschuld et al., 2012; Wolf et al., 2008).

### Perturbation of whole-brain dynamics through striatum

To estimate the baseline model, we first fitted the model for GEC of each subject, while fixing local bifurcation parameters the same across brain areas (Figure 2B). After estimating the baseline models, based on the results obtained from intrinsic ignition framework, we re-optimized the model by allowing the local bifurcation parameters of caudate and putamen to vary. Here we included putamen due to the clinical relevance of the area and that the intrinsic ignition in this area was marginally insignificant (Supplementary Materials). After re-optimizing the models, we performed in-silico experiments involving perturbation of the local bifurcation parameters of these areas (Figure 2C). The first computational experiment involved disrupting the dynamics of caudate and putamen in the model in healthy controls and measuring whether this can reproduce the FC alterations observed across HD-ISS stages. In the second computational experiment we measured how well we can recover the function of caudate and putamen in-silico by driving the activity towards healthy-like regime.

### Fitting bifurcation parameters to model disruption in striatal activity

To propose a mechanistic model, we re-optimized the model for bifurcation parameters (global, caudate and putamen) on top of the estimated EC matrices. We optimized the model for two different parameters: A global bifurcation parameter that is the same across all areas except that of caudate bifurcation parameters or putamen. First, we performed model fitting using the average GEC and FC. We found that, for the average model, global bifurcation parameters were all distributed around the value −0.3 for both HC and HD across all HD-ISS stages (Figure 3A). In contrast, caudate and putamen bifurcation parameters substantially varied between groups (Figure 3B-C). For both caudate and putamen, the bifurcation parameters were similar between HCs and Stage 0 individuals, and they were closer to 0 than the rest of the brain. These values were reduced in Stage 1 and Stage 2 to a level that they are slightly lower relative to the rest of the brain. However, in Stage 3, the bifurcation parameter, especially in caudate, was dramatically reduced.

**Figure 3.**
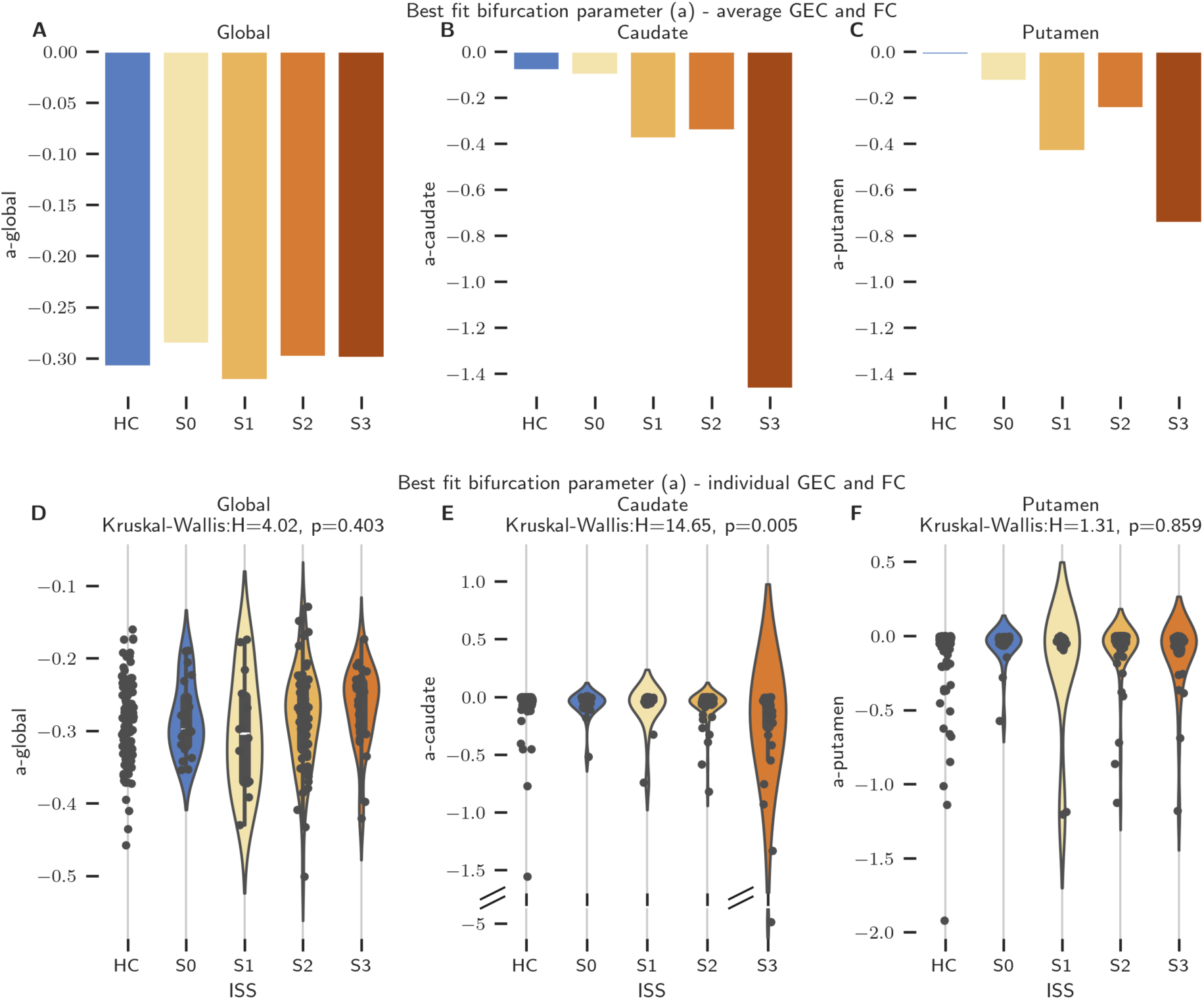
**A-C** The average model fit for bifurcation parameters (a) of caudate **(B)**, putamen **(C)** and the rest of the brain (global)**(A)**. **D-F** The individual model fit for bifurcation parameters (a) of caudate **(E)**, putamen **(F)** and the rest of the brain (global)**(D)**. The differences between means are compared using Kruskal-Wallis test.

To be able to study whether these findings can be verified statistically, we repeated the model fitting for each individual (Figure 3D-F). We found similar results, such that the global bifurcation parameters were best fit around a = −0.3, and there was no statistical difference across groups (Kruskal-Wallis test H = 4.02, p = 0.403). In contrast, we found that the bifurcation parameters of caudate were significantly different across groups (Kruskal-Wallis test H = 14.65, p = 0.005). However, we could not observe any statistical difference in bifurcation parameters of the putamen (Kruskal-Wallis test H = 1.31, p = 0.859). These results confirmed that the intrinsic dynamics of caudate are impaired in HD, especially during Stage 3 as demonstrated by intrinsic ignition framework. Although the disruption of putamen initially seemed different across HD-ISS stages from the average EC and FC matrices, this result was not supported statistically.

### In-Silico perturbation of striatum in HC participants towards HD-like dynamics

#### Average subjects

Due to the high observational noise in the data, first we performed the perturbation paradigm in average data as a proof of principle. To achieve this, we first calculated the average empirical differences in FC across HCs and HD-ISS groups. Thereafter, we established a baseline model based on average HC ECs and computed the corresponding FC, we perturbed bifurcation parameters of each area in the baseline model and recomputed the FC. Finally, we calculated how much variance (quantified by r^2^) of the empirically observed FC difference can be explained by the model perturbations.

We first focused on how much variance is explained when we perturbed caudate and putamen. As a reference, we used both the values obtained from hippocampus (like a sham stimulation), which is not known to be impaired in HD, and the average values from the rest of the brain. We found that both caudate and putamen explains a substantial amount of variance in FC differences compared to hippocampus and the rest of the brain (Figure 4A). In particular, this quantity reached up to 12% in Stage 3 caudate. Furthermore, perturbation of no other area was comparable to those of caudate perturbations in Stage 2 and Stage 3.

**Figure 4.**
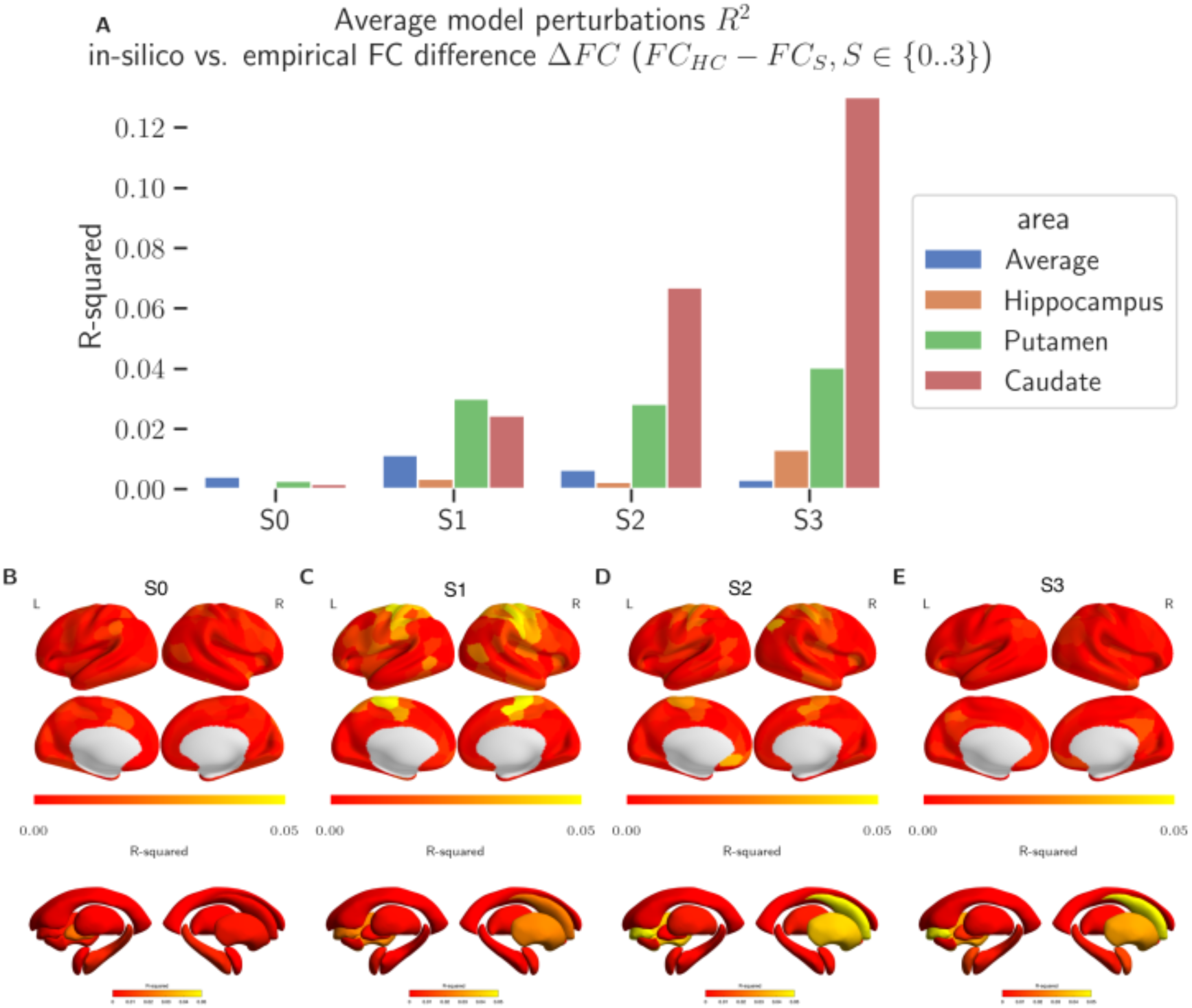
**A** In-silico perturbations of caudate and putamen in HC individuals (calculated by average EC and FC). The results indicate explained variance between empirically observed change in FC and that produced after perturbing the baseline model. The main targets for the perturbation were caudate and putamen. We show hippocampus and average across other brain regions as reference. In-silico perturbations of caudate and putamen in HC individuals (calculated by average EC and FC) for the whole-brain. **B-E** Same results for the whole-brain. The colorbars shows r-squared between empirical change in FC and those obtained in-silico.

When the perturbations are extended over all brain areas, the results clearly showed a progressive disruption in striatum (caudate and putamen) explains substantial amount of the FC differences across ISS stages (Figure 4B-E). Another surprising result that appears in the whole-brain analysis is that in STAGE 1 and STAGE 2 the perturbations in somatomotor areas also explain, albeit moderately r^2^ ∼ 4%, the FC alterations (Figure 4C-D).

#### Individual participants

After establishing the effects of perturbations of caudate and putamen in average HC FCs, we repeated the analyses in individuals. We calculated the correlation between empirical HC vs. ISS average FC differences and before/after perturbation of caudate/putamen of individual models. Note that in this case we did not use r^2^ to be able to distinguish whenever the correlation has negative values. Although the values were lower than those in the average data, there was a clear increasing trend of correlations across ISS stages (Figure 5A,C). The progression slopes were mostly close to 1 for almost all individuals as characterized by Pearson’s correlation coefficient (Figure 5B,D).

**Figure 5.**
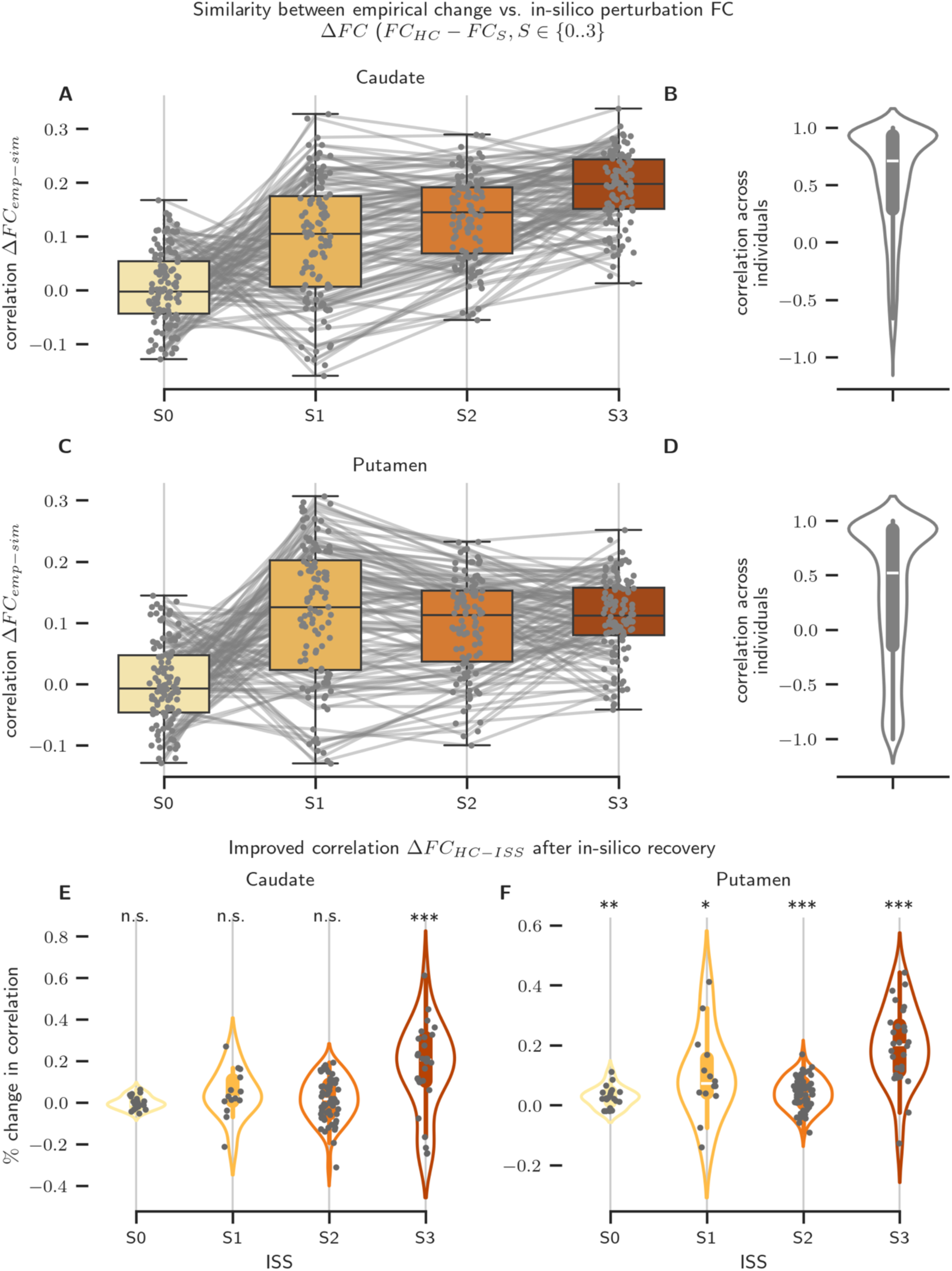
**A-B** In-silico perturbations and correlations across individuals of caudate (**A-B**) and putamen (**C-D**) in HC individuals (calculated by individual EC and FC). The values in panel A and C indicate the similarity between the FC differences across HD and HC versus simulated differences before and after perturbation. The distribution of the correlations of the similarity values along HD-ISS stages for each individual (B for caudate and D for putamen). Strong positive correlations indicate the similarity increases with disease progression as measured by ISS. **E-F** In-silico perturbations showing the recovery of FC in HD individuals after moving bifurcation parameters towards healthy dynamics for caudate (**E**) and putamen (**F**). Each dot indicates the percentage change in correlation between model and empirical FC (of HC) for a single HD patient. The distributions were tested if they are larger than 0 using one-sided T-test.

In summary, both average and individual perturbations of caudate and putamen in HCs confirmed that progressive disruptions in these areas can explain substantial variance in FC alterations. Furthermore, the model predicted that the impairment in putamen starts approximately the same time with those in caudate, it stays stable across ISS stages, whereas the impairments in caudate progressively develops across stages, hence reaching statistical significance at the later stages of the disease.

#### In-silico recovery of striatum dynamics in HD individuals

After showing that the HCs can exhibit similar FC alteration patterns to those in HD patients if caudate and putamen activity is perturbed, we studied the effects of recovering the activity of these areas in HD patients. Here we adopted a slightly different approach. After calculating the baseline model of each individual HD patient, we calculated how similar is the model FC to the empirical FC of HCs. Then we recovered the bifurcation parameters by making them halfway closer to 0 (as hypothesized that it is the case in HCs) and calculated how much the similarity between model FC and empirical FC of HCs has been improved (Figure 5E,F). This computational experiment was intended to emulate the potential effects of therapeutic stimulation of these areas. The results showed that the model was able to improve the FC very slightly in most of the cases (<1%). Interestingly, the perturbation of caudate significantly improved the FC (i.e. larger than 0) only in STAGE 3 individuals (one sided t-test T = 4.25, p < 0.001)(Figure 5E). In contrast, the perturbation of putamen lead to improved FC for all the ISS stages (one sided t-test; STAGE 0; T = 3.61, p < 0.001, STAGE 1; T = 2.67, p < 0.001, STAGE 2; T = 5.73, p < 0.001, STAGE 3; T = 8.20, p < 0.001)(Figure 5F).

## Discussion

Our study explored a theoretical framework based on intrinsic ignition to examine the effect of each individual brain region on whole-brain dynamics across different HD progression groups. First, we identified the pathological influence of the striatum, especially the caudate, on the rest of the brain. Building on this finding, we proposed a large-scale modeling framework that quantified how much of the variation in FC could be explained by activity disruption in the striatal areas. In this framework, the activity of each region varied between oscillatory and noisy dynamics depending on the local bifurcation parameter in the model. Healthy dynamics, for example, are characterized by a bifurcation parameter near its critical value, allowing a transition between different regimes. The more negative the bifurcation parameter, the more noise-driven the dynamics, mimicking a pathological state. By adjusting this bifurcation parameter, we demonstrate that virtual perturbation of the caudate and putamen in healthy individuals results in whole-brain dynamics increasingly similar to that shown in different HD-ISS stages, depending on the perturbation magnitude. Importantly, we could also replicate the opposite, where perturbing the activity of the caudate and putamen in HD-ISS stage groups could partially restore whole-brain activity to resemble healthy activity. To our knowledge, our study is the first to propose specific interventional mechanisms in HD based on in-silico models of fMRI data.

To understand the importance of our findings, we need to consider the broader context of recent HD research. First, the research community has started to change how they define and stage Huntington’s disease in its progression. Evidence of this change is the development and adoption of the HD-ISS, which incorporates the biological degeneration seen in caudate and putamen volume as an early indicator before clinical signs appear. Recognizing early degeneration has shifted research focus toward earlier HD stages, with a growing emphasis on halting or slowing biological degeneration as early as possible. Second, recent clinical trials like GENERATION-HD and PIVOT-HD have shown results indicating that treatments may be more effective when given to individuals in these early stages ((McColgan et al., 2023), PTC Therapeutics, 2025 X). As interest increasingly turns to earlier stages of the disease, there is a rising need to develop early biomarkers for HD.

Striatal atrophy has long been the key biomarker of HD progression and is also found to occur decades before the clinical motor dysfunction appears (Mena et al., 2023). While it makes sense that functional anomalies happen around the same time as structural degeneration, previous studies have shown that early structural degeneration does not match the functional changes seen in fMRI (Minkova et al., 2016; Müller et al., 2016). For example, differences in functional connectivity could not be predicted by voxel-based morphometry (VBM) changes (Müller et al., 2016) or white-matter alterations (Wolf et al., 2014). Our data suggest that abnormal FC patterns accumulate during HD progression, including Stage 1 and Stage 2, but they do not reach statistical significance until the latest stage of the disease, Stage 3. Additionally, our in-silico approach revealed that model-based disruptions in somatomotor areas partially explain the variation in whole-brain FC during Stage 1 and Stage 2. By design, the mechanisms behind these disruptions are independent from those of striatal activity.

Furthermore, we did not observe significant differences in intrinsic ignition within somatomotor areas between the stage groups in the empirical data, suggesting that the model may be more sensitive in detecting such anomalies. Additionally, abnormal connectivity in somatomotor regions peaks early on, while striatal dysconnectivity worsens at later stages. This early abnormal activity in the somatosensory cortex matches previous research showing anomalies in this region, specifically hypoconnectivity initially and hyperconnectivity later in HD (Pini et al., 2020). Because the model offers a mechanistic approach to generate altered dynamics in each brain area, our findings imply that changes in striatal dynamics do not directly cause somatosensory hypoconnectivity but instead stem from modifications in the dynamics of somatomotor areas themselves. However, understanding how this somatomotor change relates to striatal dysfunction will require further investigation, which is beyond this study’s scope. Although the model’s result matrices clearly showed differences across HD-ISS stages and identified putamen as a potential target based on average EC and FC, these results were less clear at the individual level. It is important to note the key difference between average and individual models; in the average model fitting approach, we combined the connectivity matrices of all individuals within a group to ensure that the observed effects are due to the model perturbation, and using individual connectivity estimates could lower the model fit and increase noise in the results. Using group-averaged connectivity reduces noise from systematic biases in structural connectivity estimates. This practice is common in large-scale modeling, and fitting individual-level parameters remains a significant challenge in the field (Murray et al., 2018). Furthermore, since the estimated GECs accounted for some effects caused by impairments in the caudate and putamen, these effects might not be captured by bifurcation parameters. The results could have been greatly improved by using higher temporal resolution and longer rs-fMRI data collection, along with structural connectivity (SC) estimates obtained from the same individuals.

This study identified specific effects that warrant further investigation in future research. For example, disruptions in the caudate are more progressive, with a sharper loss of function in Stage 3, while disruptions in the putamen are more moderate, beginning as early as Stage 1 and remaining stable as the disease progresses. Additionally, the model predicts that disruptions in somatomotor areas occur during middle stages of the disease (i.e., Stage 1 and Stage 2) and then return to normal at Stage 3. Lastly, our modeling framework indicates that the putamen may be a more effective target than the caudate for early-stage therapeutic interventions.

It is important to acknowledge the relevance of our findings within the context of this study’s limitations. The most significant limitation we faced was the need to concatenate or average group data to detect key differences between groups, likely due to lack of robust data. Although TRACKOn-HD is one of the highest quality and most comprehensive HD datasets in terms of structural imaging and phenotypic data acquisition, its scan duration of 8.25 minutes and sampling frequency (TR = 3s) fall below the recommended standards for optimal resting state data collection. Several studies have demonstrated that increasing fMRI scan durations to over 13 minutes up to 25 minutes, and improving temporal resolution, can greatly enhance sensitivity and the stability of results (Birn et al., 2013; Caeyenberghs et al., 2024; see review by Noble et al., 2019). After concatenating each subject’s longitudinal data, we created individual time series up to 25 minutes long, which indeed increased analysis sensitivity.

Additionally, the stage groups examined were imbalanced, and the sample sizes were sometimes suboptimal. For example, one group had N = 13, which is below the recommended minimum of N approximately 40 (Ma et al., 2024). This discrepancy occurred because the TRACKOn-HD study was designed with different grouping criteria, and we had to use this dataset to explore HD progression using the newer HD-ISS criteria. Another important point is that intrinsic ignition values in many brain regions were significantly correlated with in-scanner motion (i.e., framewise displacement; see Supplementary Material). Although there was no statistical difference in motion between groups, this confounding factor likely added variance to the data, further limiting the ability to model and compare specific brain regions. Notably, our findings from in-silico perturbations are unaffected by head motion since they involve virtual perturbation of a single parameter rather than direct group comparisons. Lastly, a potential limitation that is difficult to address is the partial volume effect, especially as structures like the caudate and putamen shrink. Smaller structures are more likely to be covered by voxels that include neighboring tissues such as ventricles and white matter. As the striatum atrophies, the BOLD signal may become contaminated by signals from surrounding regions, reducing the number of voxels fully *within* the structure. Consequently, the fMRI activity comparison in caudate and putamen between stage groups may contain more noise due to atrophy rather than genuine fMRI signal changes. Although this issue could influence direct group comparisons, we also used perturbations of healthy individuals to simulate activity, which gradually resembled HD progression. The similarity between these in-silico perturbations and empirical BOLD measurements suggests our findings are not primarily driven by partial voluming artifacts.

In conclusion, we adopted an in-silico approach to emulate noninvasive brain stimulation in HD. Our results suggested that it is possible to recover FC, albeit moderately, by tuning the local bifurcation parameter of caudate and putamen, hence pushing them towards the edge of criticality, that is the proximity of a dynamical system to a critical point in which the system exhibits transition between different regimes (Deco et al., 2017b; Deco and Jirsa, 2012). Furthermore, our model predicted that although disruptions of the caudate are more influential on overall brain activity, maintaining healthy putamen activity might be a more effective therapeutic approach in earlier stages. Real stimulation of striatal activity in healthy individuals has also been shown to increase improvement in motor performance when the putamen, rather than the caudate, responds to the stimulation (Wessel et al., 2023). Despite our encouraging in-silico results, this study only provides a mechanistic interpretation of the directions the interventions should follow; it is not possible to determine a frequency-specific, rigorous stimulation protocol based solely on the fMRI data at hand. Altogether, this study demonstrates that computational modeling is a valuable tool understand the underlying patterns behind the observed data and propose causal mechanisms for empirical testing.

## Notes

### Competing Interest Statement

The authors have declared no competing interest.

### Funding

- CHDI Foundation: A-17884

